# Whole genome sequencing *Mycobacterium tuberculosis* directly from sputum identifies more genetic diversity than sequencing from culture

**DOI:** 10.1101/446849

**Authors:** Camus Nimmo, Liam P. Shaw, Ronan Doyle, Rachel Williams, Kayleen Brien, Carrie Burgess, Judith Breuer, Francois Balloux, Alexander S. Pym

## Abstract

**Background:** Repeated culture reduces within-sample *Mycobacterium tuberculosis* genetic diversity due to selection of clones suited to growth in culture and/or random loss of lineages, but it is not known to what extent omitting the culture step altogether alters genetic diversity. We compared *M. tuberculosis* whole genome sequences generated from 33 paired clinical samples using two methods. In one method DNA was extracted directly from sputum then enriched with custom-designed SureSelect (Agilent) oligonucleotide baits and in the other it was extracted from mycobacterial growth indicator tube (MGIT) culture.

**Results:** DNA directly sequenced from sputum showed significantly more within-sample diversity than that from MGIT culture (median 5.0 vs 4.5 heterozygous alleles per sample, p=0.04). Resistance associated variants present as HAs occurred in four patients, and in two cases may provide a genotypic explanation for phenotypic resistance.

**Conclusions:** Culture-free *M. tuberculosis* whole genome sequencing detects more within-sample diversity than a leading culture-based method and may allow detection of mycobacteria that are not actively replicating.

## Background

International efforts to reduce tuberculosis (TB) infections and mortality over the last two decades have only been partially successful. In 2017, 10 million people developed TB and it has overtaken HIV as the infectious disease responsible for the most deaths worldwide(1, 2). Drug resistance is a major concern with a steady rise in the number of reported cases globally and rapid increases in some areas(1). Patients with *Mycobacterium tuberculosis* resistant to the first line drugs rifampicin and isoniazid are classed as having multidrug-resistant (MDR) TB and usually treated with a standardised second line drug regimen for at least nine months, which is also used for rifampicin monoresistance(3, 4). With the emergence of resistance to fluoroquinolones and aminoglycosides (extensively drug-resistant [XDR] TB) there is an increasing need for individualised therapy based on drug susceptibility testing (DST). Individualised therapy ensures patients are treated with sufficient active drugs which can prevent selection of additional resistance, improve treatment outcomes and reduce duration of infectiousness(5–8).

Traditionally, phenotypic culture-based DST was used to identify drug resistance but this is being replaced by rapid genetic tests that detect specific drug resistance-conferring mutations. Next generation whole genome sequencing (WGS) of *M. tuberculosis* is being increasingly used in research and clinical settings to comprehensively identify all drug resistance associated mutations(9). *M. tuberculosis* has a conserved genome with little genetic diversity between strains and no evidence of horizontal gene transfer(10), but more detailed analysis of individual patient samples with WGS has identified genetically separate bacterial subpopulations in sequential sputum samples(11–16) and across different anatomical sites(17). This within-patient diversity can occur as a result of mixed infection with genetically distinct strains or within-host evolution of a single infecting strain(18).

Bacterial subpopulations can be detected in clinical samples after sequencing reads are mapped to a reference genome where multiple base calls are detected at a single genomic site. These heterozygous alleles (HAs) at sites associated with drug resistance (resistance associated variants, RAVs) may reflect heteroresistance, where a fraction of the total bacterial population is drug susceptible while the remainder is resistant(19). Identification of genetic diversity within clinical samples may improve detection of RAVs over currently available rapid genetic tests(19) and can be achieved with freely available WGS analysis toolkits(20–22). Identifying RAVs could improve individualised therapy, prevent acquired resistance(12), and give insight into bacterial adaptation to the host.

*M. tuberculosis* WGS is usually performed on fresh or stored frozen cultured isolates to obtain sufficient purified mycobacterial DNA(23, 24). However, the culture process can change the population structure from that of the original sample due to genetic drift (random loss of lineages) and/or the selection of subpopulations more suited to growth in culture(25–27). Repeated subculture leads to loss of genetic diversity and heteroresistance(28). Additionally, in the normal course of *M. tuberculosis* infection, some bacteria exist as viable non-culturable persister organisms that are hypothesised to cause the high relapse rate seen following treatment of insufficient duration. Although these organisms may be identified in sputum by techniques such as reporter phages or culture with resuscitation promoting factors(29, 30) they are likely to be missed by any sequencing method reliant on standard culture.

WGS directly from sputum without enrichment is challenging(23). It has recently been improved by depleting human DNA during DNA extraction(31). We have previously reported the use of oligonucleotide enrichment technology SureSelect (Agilent, CA, USA) to sequence *M. tuberculosis* DNA directly from sputum(32) and demonstrated its utility in determining a rapid genetic drug resistance profile(33, 34).

It remains unclear to what extent WGS of cultured *M. tuberculosis* samples underestimates the genetic diversity of the population in sputum samples. One previous study of 16 patients did not identify increased genetic diversity in *M. tuberculosis* DNA sequenced directly from sputum compared to DNA from culture(31), whereas another study of mostly drug susceptible patients showed sequencing directly from sputum identified a slight excess of HAs relative to culture(33). Here we reanalyse heterozygous alleles (HAs) for the 12 available paired sequences with >60-fold mean genome coverage from that study(33) in addition to 21 newly collected samples from patients with MDR-TB and further explore the genomic location of the additional diversity identified.

## Results

### Patient Characteristics and Drug Susceptibility Testing

Whole genome sequences were obtained for 33 patients from both mycobacterial growth indicator tube (MGIT) culture and direct sputum sequencing. The patients were predominantly of black African ethnicity (83%) and 50% were HIV positive. First line phenotypic drug susceptibility testing (DST) results identified 20 patients with MDR-TB and one with rifampicin monoresistance. In addition there were two isoniazid monoresistant patients and ethambutol resistance was detected in 7 patients. Second-line phenotypic DST was performed for patients with rifampicin-resistant or MDR-TB and identified one case of kanamycin resistance (Table 1).

**Table 1.**
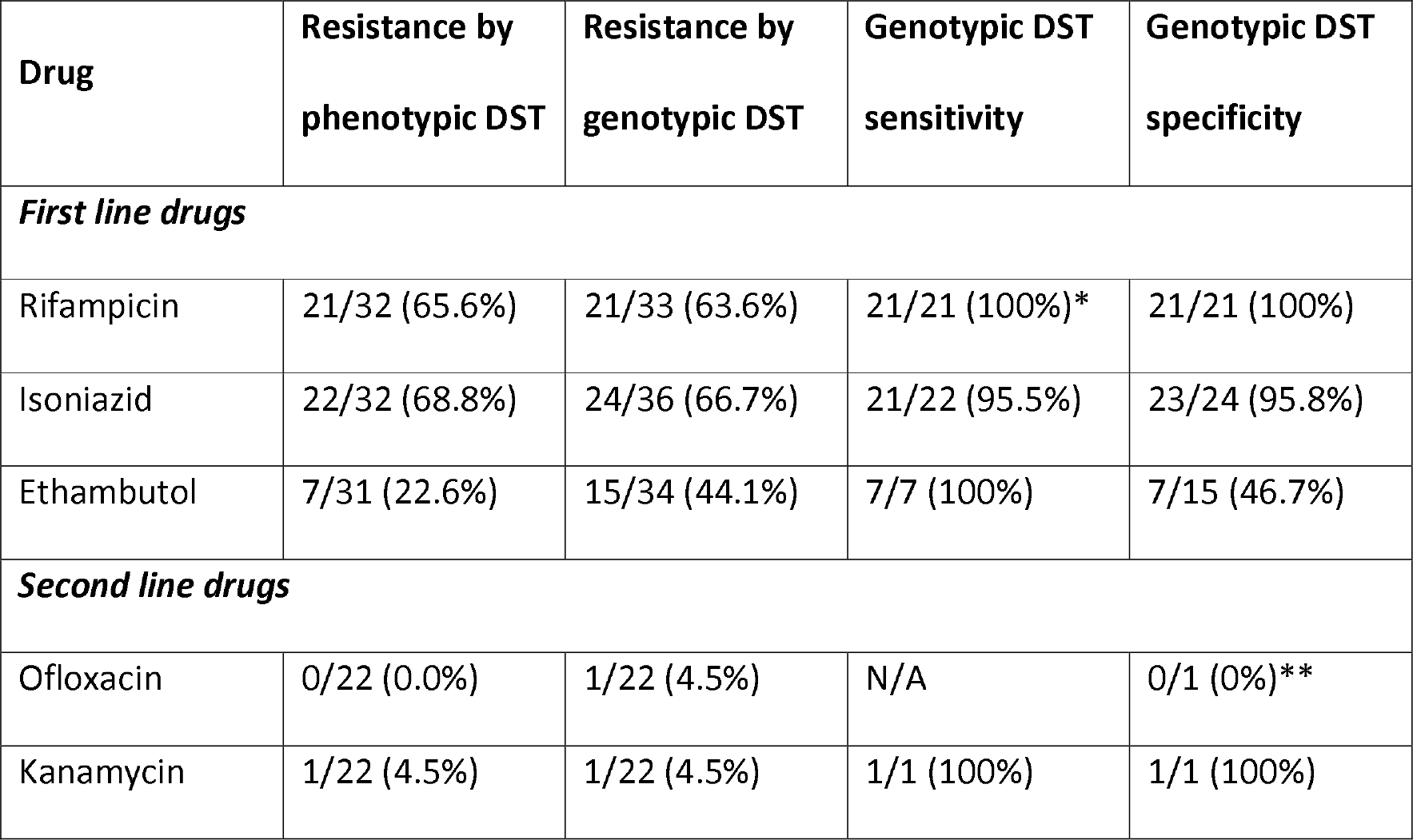
Phenotypic and genotypic drug susceptibility testing (DST) results and sensitivity and specificity of genotypic DST relative to phenotypic DST. Phenotypic DST available for first line drugs for 32 of the 33 patients, and for second line drugs for 22 patients who demonstrated rifampicin drug resistance. *In one directly-sequenced sputum samples rifampicin RAVs were missed due to low coverage, although they were identified in the corresponding MGIT sample. **This sample had <1% of colonies grow in the presence of ofloxacin, so is categorised as sensitive but may have low-level or heteroresistance to fluoroquinolones (see main text).

| Drug | Resistance by phenotypic DST | Resistance by genotypic DST | Genotypic DST sensitivity | Genotypic DST specificity |
| --- | --- | --- | --- | --- |
| ***First line drugs*** |  |  |  |  |
| Rifampicin | 21/32 (65.6%) | 21/33 (63.6%) | 21/21 (100%)* | 21/21 (100%) |
| Isoniazid | 22/32 (68.8%) | 24/36 (66.7%) | 21/22 (95.5%) | 23/24 (95.8%) |
| Ethambutol | 7/31 (22.6%) | 15/34 (44.1%) | 7/7 (100%) | 7/15 (46.7%) |
| ***Second line drugs*** |  |  |  |  |
| Ofloxacin | 0/22 (0.0%) | 1/22 (4.5%) | N/A | 0/1 (0%)** |
| Kanamycin | 1/22 (4.5%) | 1/22 (4.5%) | 1/1 (100%) | 1/1 (100%) |

All samples had mean genome coverage of 60x or above with at least 85% of the genome covered at 20x (Supplementary Material: Table 1). We observed greater mean coverage depth in sputum-derived sequences than MGIT sequences (median 173.7 vs 142.4, p=0.03, Supplementary Material: Table 1), and so mapped reads were randomly downsampled to give equal mean coverage depth in each pair. A genotypic susceptibility profile was determined by evaluating MGIT WGS for consensus-level RAVs using a modified version of publicly available lists(22, 35). Genotypic RAVs predicted all rifampicin phenotypic resistance and >95% of isoniazid phenotypic resistance. Ethambutol genotypic RAVs were poorly predictive of phenotypic resistance in line with findings from other studies(36) (Table 1). The patient with kanamycin phenotypic resistance was correctly identified by an *rrs* a1401g RAV. No full phenotypic fluoroquinolone phenotypic resistance was identified, but several colonies from patient F1013 did grow in the presence of ofloxacin (although not enough to be classified as resistant). The consensus sequences from this patient harboured a *gyrB* E501D mutation which is believed to confer resistance to moxifloxacin but not other fluoroquinolones, which may explain the borderline phenotypic DST result(37).

### Genetic Diversity

To compare consensus sequences from sputum and MGIT, a WGS consensus sequence-level maximum likelihood phylogenetic tree was constructed (Supplementary Material: Figure 1). As expected, all paired sequences were closely related, with a median difference of 0.0 (range 0-1) single nucleotide polymorphisms (SNPs). Samples from patients F1066 and F1067 were closely related with only one consensus-level SNP separating all four consensus sequences. There was no obvious epidemiological link between these patients (although this study was not designed to collect comprehensive epidemiological information) and they lived 20km apart in Durban. However, both patients were admitted contemporaneously to an MDR treatment facility and sampled on the same day. DNA extraction and sequencing occurred on different runs. Therefore the close genetic linkage may represent direct transmission within a hospital setting, a community transmission chain or an unlikely cross-contamination during sample collection.

Having established congruence between sputum and MGIT sequences at the consensus level we then compared genetic diversity by DNA source. We first defined a threshold for calling variants present as heterozygous alleles (HAs) in our entire dataset by using a range of minimum read count frequencies as described in the methods (Figure 1). Below a minimum of three supporting reads there was an exponential increase in the number of HAs identified, which may be indicative of the inclusion of sequencing errors. To reduce this risk, we used a threshold of a minimum of four supporting reads.

**Figure 1.**
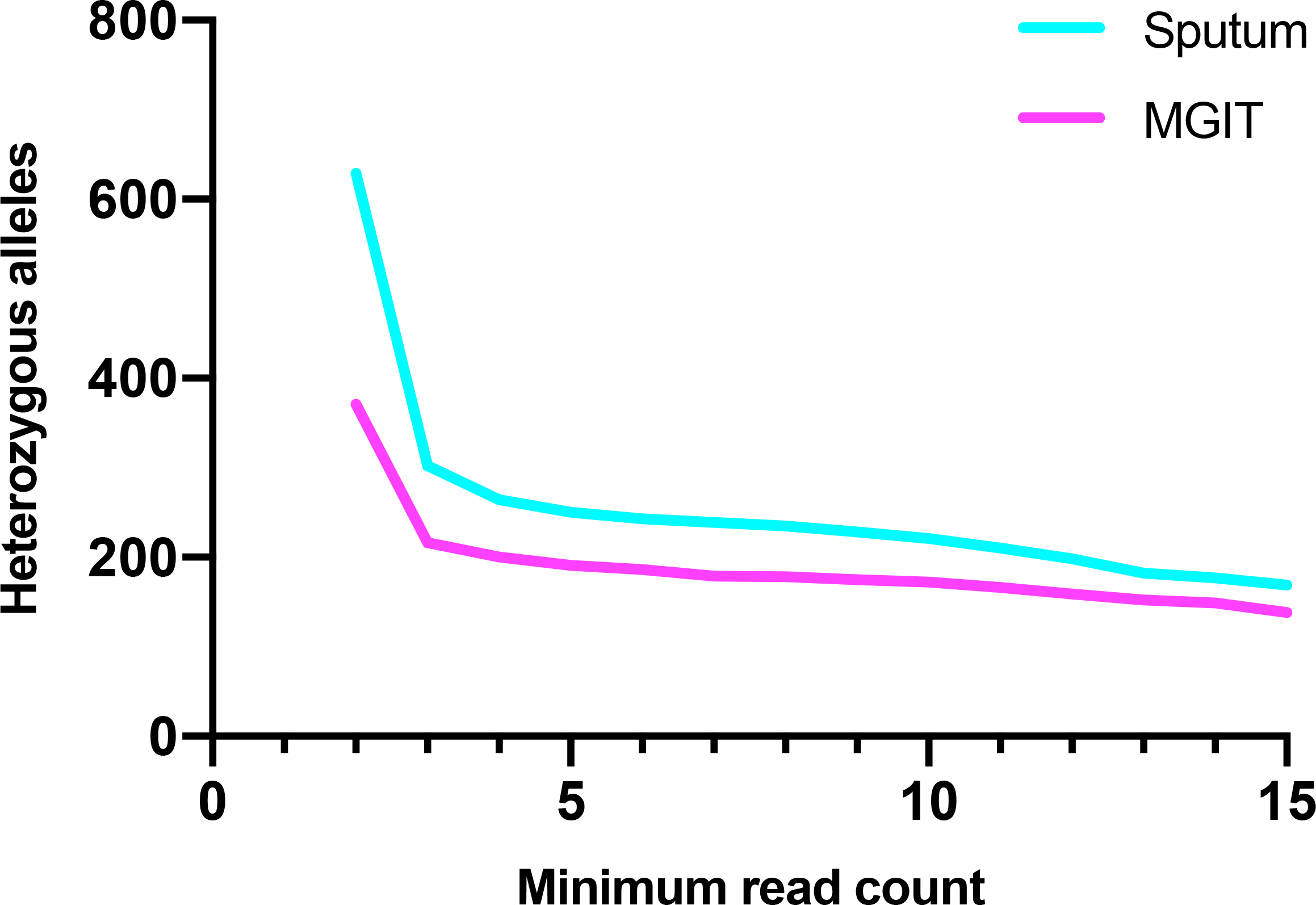
Variation in total number of heterozygous alleles (HAs) identified across all 36 patients in sequences generated from sputum and MGIT depending on minimum supporting read count threshold.

Genetic diversity may occur because of within-host evolution or mixed infection. To identify mixed infection we used a SNP-based barcode(38) to scan all HAs for a panel of 413 robust phylogenetic SNPs that can resolve *M. tuberculosis* into one of seven lineages and 55 sub-lineages. We found three phylogenetic SNPs among the HAs. In all cases the heterozygous phylogenetic SNP originated from the same sublineage as other SNPs present at 100% frequency, and there were no cases of HAs indicating the presence of more than one lineage or sublineage. We screened for mixed infection with the same sublineage by screening samples by HA frequency and then using Bayseian model based clustering in samples with ≥10 HAs as described previously(39). This identified mixed infection in the sputum sample from patient F1096, which had 261 heterozygous alleles, greater than ten times that in any other sample. This patient was therefore excluded from further analyses.

As a first step to comparing diversity between sputum and MGIT sequenced samples we looked at the location of genetic diversity within the *M. tuberculosis* genome. HAs were widely dispersed across the genome at similar sites in both sputum and MGIT samples. The genes with the greatest density of HAs are shown in Table 2.

**Table 2.** Genes with ≥2 heterozygous alleles (HAs) across all sputum samples, ordered by greatest number of HAs per base.

| Gene | Heterozygous alleles<br>per base |  | Total number of<br>heterozygous alleles |  | Functional category |
| --- | --- | --- | --- | --- | --- |
|  | Sputum | MGIT | Sputum | MGIT |  |
| *rv1319c* | 0.021 | 0.021 | 33 | 33 | Metabolism and<br>respiration |
| *rrs* | 0.016 | 0.000 | 25 | 0 | 16S ribosomal RNA |
| *rrl* | 0.006 | 0.000 | 19 | 0 | 23S ribosomal RNA |
| *ppsA* | 0.003 | 0.001 | 15 | 4 | Lipid metabolism |
| *rv2082* | 0.006 | 0.006 | 13 | 14 | Unknown function |
| *accE5* | 0.006 | 0.000 | 3 | 0 | Lipid metabolism |
| *lppB* | 0.005 | 0.005 | 3 | 3 | Probable surface<br>lipoprotein |
| *pks12* | 0.000 | 0.001 | 3 | 10 | Lipid metabolism |
| *rv2319c* | 0.003 | 0.005 | 3 | 4 | Stress protein |
| *lppA* | 0.003 | 0.002 | 2 | 1 | Probable surface<br>lipoprotein |
| *rpoC* | 0.001 | 0.001 | 2 | 3 | RNA polymerase beta'<br>subunit |
| *rv3888c* | 0.002 | 0.001 | 2 | 1 | Probable membrane<br>protein |
| *vapC25* | 0.005 | 0.000 | 2 | 0 | Possible toxin |
| *vapC31* | 0.005 | 0.002 | 2 | 1 | Possible toxin |

Notably, genetic diversity was found in the ribosomal RNA (rRNA) genes (*rrs* and *rrl*) uniquely in sputum samples, compared to other genes where distribution of diversity between MGIT and sputum was more balanced. As rRNA contains regions that are highly conserved across bacteria(40), we considered the possibility that SureSelect baits targeting rRNA genes were capturing both *M. tuberculosis* and other bacterial species. To evaluate this, metagenomic taxonomic assignment was performed on all reads by sampling reads that were not assigned to *M. tuberculosis* (i.e. presumed contaminants from other bacteria). We then performed a BLAST search against the most diverse genes listed in Table 2 which indicated that a sizeable proportion of non-*M. tuberculosis* reads from directly sequenced sputum had a BLAST hit of at least 30 bases to *M. tuberculosis rrs* and *rrl* genes that encode rRNA (330 BLAST hits from sputum sequences vs 4 BLAST hits from MGIT sequences, median 8.5% vs 0.0%, p<0.01, Supplementary Material: Figure 2). There were no BLAST hits against any of the other genes with ≥2 sputum HAs apart from *rpoC*, for which there were 3 BLAST hits from sputum sequences but none from MGIT sequences (median 0.0% for both sputum and MGIT sequences), indicating that this issue appears largely specific to rRNA. To determine if contaminating reads were contributing to HAs identified in intergenic regions, we repeated this analysis for all intergenic regions with ≥2 sputum HAs (Supplementary Material: Table 2). There were no BLAST hits to any of these regions, suggesting that this is not the case. The taxonomic assignment of these contaminating reads were typical of genera composing the oral flora, with a high representation of *Actinomyces*, *Fusobacterium*, *Prevotella*, and *Streptococcus* (Supplementary Material: Figure 3).

This supported the hypothesis that the baits may enrich rRNA from other organisms so rRNA genes were excluded from further analysis. The difference in diversity between sputum and MGIT sequences can be explained by the selective nature of MGIT media which will enrich *M. tuberculosis* sequences and the decontamination step used to kill non-mycobacteria prior to culture inoculation. Importantly the frequency of HAs in other highly diverse genes between sequencing strategies was more balanced (Table 2) in addition to the lack of BLAST hits of contaminating reads to these genes.

After excluding the sample with mixed infection and removing rRNA gene sequences we compared the frequency of HAs in sputum and MGIT. There were 265 HAs identified across all sputum samples compared to 200 in MGIT samples (median 5.0 vs 4.5, p=0.04, Supplementary Material: Table 1). In both sputum and MGIT samples, the majority of Has were indels, and non-synonymous mutations were more commonly frameshift than missense mutations (Table 3). The distribution of HAs by patient is shown in Figure 2.

**Table 3.** Variants identified in MGIT and sputum derived sequences from paired samples. Values given represent totals for 32 paired samples. SNP = single nucleotide polymorphism; MNP = multi-nucleotide polymorphism.

|  | **Sputum variants** | **MGIT variants** |
| --- | --- | --- |
| Total variants | 24480 | 25465 |
| Total variants present as HAs<br>(% of total variants) | 265 (1.1%) | 200 (0.8%) |
| Median HAs per sample | 5.0 | 4.5 |
| Variant type (% all HAs) |  |  |
| SNP | 217 (81.9%) | 174 (87.0%) |
| MNP | 2 (0.8%) | 0 (0.0%) |
| Insertion | 4 (1.5%) | 1 (0.5%) |
| Deletion | 24 (9.1%) | 15 (7.5%) |
| Complex | 18 (6.8%) | 10 (5.0%) |
| Coding change (% all HAs) |  |  |
| Non-synonymous (missense) | 93 (35.1%) | 77 (38.5%) |
| Non-synonymous (frameshift) | 6 (2.3%) | 7 (3.5%) |
| Synonymous | 57 (21.5%) | 57 (28.5%) |
| Intergenic | 109 (41.1%) | 59 (29.5%) |

**Figure 2.**
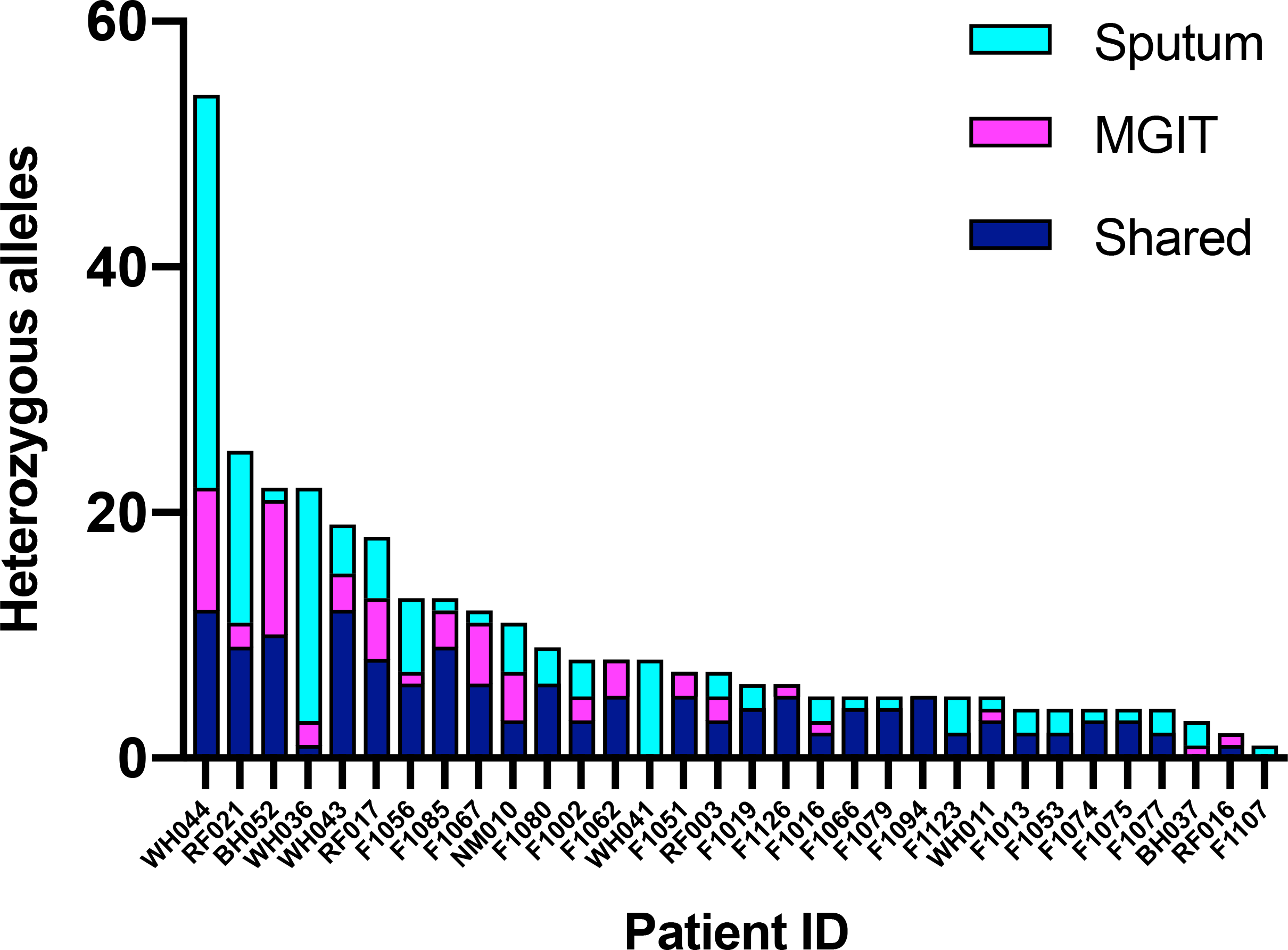
Number of heterozygous alleles (HAs) found in directly sequenced sputum only (sputum), MGIT (MGIT) only or in both samples (shared) by patient.

### Genetic diversity in drug resistance genes

HAs in drug resistance associated regions, including promoters and intergenic regions, were individually assessed. Four of the 32 patients with single strain infection had RAVs present as HAs in at least one gene, which are shown in Table 4. Patient F1002 had three compensatory mutations in *rpoC* present at HAs in both sequences. As described above, the strains from patients F1066 and F1067 were highly related with only one consensus SNP difference between all four sequences. Both had phenotypic high level isoniazid resistance with no consensus-level *katG* or *inhA* mutation, but had frameshift *katG* mutations present as HAs which have the potential to cause resistance(41). F1066 and RF021 had *Rv1979c* and *pncA* mutations respectively at low frequency in sputum only which have the potential to confer phenotypic resistance to clofazimine (*Rv1979c*) and pyrazinamide (*pncA*), although no phenotypic testing was performed for these drugs.

**Table 4.** Resistance associated variants present as heterozygous alleles (HAs).

| Patient ID | Phenotypic resistance | Mutation | Frequency (MGIT/sputum) | Description |
| --- | --- | --- | --- | --- |
| F1002 | Rifampicin | *rpoB* S450L | 100%/100% | High confidence resistance mutation |
| F1002 | Rifampicin | *rpoC* G332R(58) | 82.6%/21.7% | Putative compensatory mutations |
| F1002 | Rifampicin | *rpoC* L516P(58) | 12.7%/7.7% |  |
| F1002 | Rifampicin | *rpoC* P1040S(59) | 21.7%/12.3% |  |
| F1066 | Isoniazid (high) | *katG* N218fs | 0.0%/6.9% | Possible resistance mutations, not previously described |
| F1066 | Clofazimine – not tested | *Rv1979c* G376D | 0.0%/0.5% |  |
| F1067 | Isoniazid (high) | *katG* N218fs | 10.7%/7.6% |  |
| RF021 | Pyrazinamide – testing failed | *pncA* Q122H | 0%/2.5% |  |

## Discussion

In this study we performed whole genome sequencing using DNA from sputum and MGIT culture in paired samples from 33 patients and compared within-patient genetic diversity between methods. All paired sequences were closely related at the consensus level, and WGS predicted phenotypic drug susceptibility with over 95% sensitivity and specificity for rifampicin and isoniazid in line with published data(42).

We find that the rRNA genes have high levels of diversity in sputum samples, but believe this is due to non-mycobacterial DNA hybridising to the capture baits. This conclusion is borne out by the taxonomic assignment of reads aligning to these genes in common oral bacteria. We therefore excluded these from further analysis, and recommend others using enrichment from sputum do similarly. We find more diversity when sequencing directly from sputum with significantly more unique heterozygous alleles (HAs) than sequencing from MGIT culture (p=0.04).

The understanding of within-patient *M. tuberculosis* genetic diversity is becoming increasingly important as the detection of rare variants has been shown to improve the correlation between phenotypic and genotypic drug resistance profiles(19) and can identify emerging drug resistance(11, 12). Not including a culture step avoids the introduction of bias towards culture-adapted subpopulations and the impact of random chance and is also likely to incorporate DNA from viable non-culturable mycobacteria. A reduction in genetic diversity has previously been shown with sequential *M. tuberculosis* subculture(25, 28), but was not confirmed by a study performing WGS directly from sputum(31). However, the 16 paired sputum and MGIT samples compared by Votintseva(31) had a minimum of 5x coverage compared to a minimum 60x coverage in this study, and were likely to contain less genetic material as they were surplus clinical rather than dedicated research samples.

Two-thirds of the patients with MDR-TB had already been treated for drug susceptible-TB (DS-TB), and additional diversity in sputum samples may represent early adaptation to drug pressure. As direct sputum sequencing does not rely on live mycobacteria, DNA from recently killed *M. tuberculosis* is likely to also be sequenced, meaning that recent genomic mutations are likely to be represented as HAs.

In two patients, RAVs present as HAs provided a likely genotypic basis for otherwise unexplained phenotypic resistance. Given the small total number of resistance mutations in this study, it is not possible to draw conclusions about the frequency of heterozygous RAVs in directly sequenced sputum. However the presence of heterozygous RAVs in both MGIT and sputum sequences reinforces the biological importance of these mutations.

To reduce the risk of sample cross contamination, paired samples were extracted on different days, prepared in different sequencing libraries and sequenced on different runs. However it is not possible to completely exclude the possibility of contamination during sample collection and between different samples processed in batches. A further limitation of this study is that it can be difficult to distinguish low frequency variants from sequencing error. The SureSelect library preparation protocol for sputum sequencing incorporates more PCR cycles than that used for MGIT sequencing, which may increase the risk of error. Where possible this could be evaluated further by performing technical sequencing replicates on extracted DNA samples, although this was not possible due to insufficient surplus material and financial constraints. To reduce the risk of sequencing errors we used high read and mapping quality thresholds, and required a stringent 98% identity between sequenced reads and the reference genome. Low frequency variants of particular clinical importance could be confirmed by resequencing the same DNA samples.

## Conclusions

Directly sequencing *M. tuberculosis* from sputum is able to identify more genetic diversity than sequencing from culture. Understanding within-patient genetic diversity is important to understand bacterial adaptation to drug treatment and the acquisition of drug resistance. It also has potential to identify low frequency RAVs that may further enhance the prediction of drug resistance phenotype from genotype.

## Methods

### Patient enrolment

Adult patients presenting with a new diagnosis of sputum culture positive TB were included in the study. Patients were recruited in London, UK (n=12) and Durban, South Africa (n=21). All patients recruited in Durban were Xpert MTB/RIF (Cepheid, CA, USA) positive for rifampicin resistance. Two sputum samples were collected prior to starting the current treatment regimen, with one inoculated into mycobacterial growth indicator tube (MGIT) culture (BD, NJ, USA) and the other used for direct DNA extraction. Therefore for patients with drug susceptible-TB (DS-TB), sputum was collected prior to taking any TB therapy, while patients starting MDR-TB treatment may have already taken treatment for DS-TB if this was intiated prior to resistance results being available.

### Ethics, Consent and Permissions

All patients gave written informed consent to participate in the study. Ethical approval for the London study was granted by NHS National Research Ethics Service East Midlands–Nottingham 1 (reference 15/EM/0091). Ethical approval for the Durban study was granted by University of KwaZulu-Natal Biomedical Research Ethics Committee (reference BE022/13).

### Microbiology

MGIT samples were incubated in a BACTEC MGIT 960 (BD, NJ, USA) until flagging positive. Phenotypic DST data for London samples were those provided to treating hospitals by Public Health England. Phenotypic DST were performed using equivalent standardised methods. For Durban samples this was the solid agar proportion method (Supplementary Material: Methods) and for London samples the resistance ratio method(43).

### DNA extraction and sequencing

Positive MGIT tubes were centrifuged at 16,000g for 15 minutes and the supernatant removed. Cells were resuspended in phosphate-buffered saline before undergoing heat killing at 95°C for 1 hour followed by centrifugation at 16,000g for 15 minutes. The supernatant was removed and the sample resuspended in 1mL sterile saline (0.9% w/v). The wash step was repeated. DNA was extracted with mechanical ribolysis before purification with DiaSorin Liaison Ixt (DiaSorin, Italy) or CTAB(44). NEBNext Ultra II DNA (New England Biolabs, MA, USA) was used for DNA library preparation.

Sputum samples for direct sequencing were heat killed, centrifuged at 16,000g for 15 minutes and the supernatant was removed. DNA extraction was performed with mechanical ribolysis followed by purification using DiaSorin Liaison Ixt (DiaSorin, Italy) or DNeasy blood & tissue kit (Qiagen, Germany)(44). Target enrichment was performed using SureSelect with a custom-designed bait set covering the entire positive strand of the *M. tuberculosis* genome as described previously(33). Batches of 48 multiplexed samples were sequenced on NextSeq 500 (Illumina, CA, USA) 300-cycle paired end runs with a mid-output kit. Sequencing was performed by the Pathogen Genomics Unit at University College London in a dedicated laboratory where one sequencing run was processed at a time. All paired samples were extracted, prepared and sequenced on different days. The National Center for Biotechnology Information Sequence Read Archive (NCBI SRA) accession number for each sample is shown in Supplementary Material: Table 3.

### Read mapping

DNA sequence reads were adapter and quality trimmed then aligned to the H37Rv reference genome (GenBank accession NC_000962.3) with Trim Galore v0.4.4(45) and BBMap v38.32(46), with mapped reads stored in an output bam file. Duplicate reads were removed with Picard tools v1.130(47) MarkDuplicates and coverage statistics generated with Qualimap v2.2.1(48). For each sample pair, the bam file with greater mean genome coverage was randomly downsampled to that of the paired sample with Picard tools v1.130(47) DownsampleSam. All further analyses were performed using these downsampled bam files. Command line parameters used are specified in the Supplementary Material: Methods.

### Variant calling

Variant calling for comparison for HA counts was performed with FreeBayes v1.2(49). Variants falling in or within 50 bases of PE/PPE family genes and repeat elements were excluded using vcfinteresect in vcflib(50). For the initial analysis of genetic diversity, variants were included if supported by ≥2 reads, with ≥1 forward and reverse read, no read position bias, a minimum mapping quality of 30 and base quality of 30. The minimum supporting read threshold was then increased in a stepwise fashion from 2 to 15. Variant calling files where variants were supported ≥4 supporting reads including ≥1 forward and reverse read were used to compare HA frequency and location and to screen for mixed infection.

The phylogenetic tree was constructed by calling variants with VarScan v2.4.0(51) mpileup2cns as this is able to generate consensus-level calls at each reference sequence base. SNPs were then used to generate a sequence of equal length to the reference using a custom perl script and these sequences were combined in a multi-alignment fasta file. SNP sites were extracted from this alignment using snp-sites v2.4.1(52), and pairwise SNP differences calculated using snp-dists v0.6.3(53). Extracted SNP sites were used to generate a maximum likelihood phylogenetic tree using RaxML v8.2.12(54) which was visualised using FigTree v1.4.3.

### Identification of Mixed Infection

All samples were screened for evidence of mixed infection using described methods(39). In brief, any sample with 10 or fewer heterozygous SNPs, or between 11 and 20 heterozygous SNPs where heterozygous SNPs were ≤1.5% of all SNPs was classified as not mixed. For other samples, the Baysian mixture model analysis(39) was used where samples with a Bayesian information criterion value >20 for presence of more than one strain were assumed to be mixed.

### Metagenomic assignment

Sequencing reads were classified using Kraken v0.10.6(55) against a custom Kraken database previously constructed from all available RefSeq genomes for bacteria, archaea, viruses, protozoa, and fungi, as well as all RefSeq plasmids (as of September 19^th^ 2017) and three human genome reference sequences(56). The size of the final database after shrinking was 193 Gb, covering 38,190 distinct NCBI taxonomic IDs.

To assess the proportion of contaminating reads that could generate spurious diversity when mapped to *M. tuberculosis* ribosomal genes, we randomly subsampled 100 reads taxonomically assigned as non-*M. tuberculosis* and performed a BLAST search with blastn v2.2.28(57) against each gene as described from the H37Rv reference genome. We only analysed hits of at least 30 bases.

### Statistics

Statistical analyses were performed with Prism v8.0 (Graphpad, CA, USA). Mean coverage depth statistics, number of HAs and BLAST hits of contaminating reads in paired samples were compared using a two-tailed Wilcoxon matched-pairs signed rank test.

## Abbreviations

DST: drug susceptibility testing
DS-TB: drug susceptible-tuberculosis
HA: heterozygous allele
MDR-TB: multidrug resistant-tuberculosis
MGIT: mycobacterial growth indicator tube
RAV: Resistance associated variant
rRNA: ribosomal RNA
SNP: single nucleotide polymorphism
TB: tubersulosis
WGS: whole genome sequencing

## Declarations

### Consent for publication

Not applicable

### Availability of data and materials

Original fastq files are available at NCBI Sequence Read Archive with BioProject reference PRJNA486713: https://www.ncbi.nlm.nih.gov/bioproject/PRJNA486713/

### Authors’ contributions

Study conception: JB, ASP

Data collection: CB, KB

Analysis and interpretation: CN, LPS, RD, RW

Drafting of manuscript: CN, LPS

Revision of manuscript: FB, JB, ASP

Final approval of manuscript: CN, LPS, RD, RW, KB, CB, JB, FB, ASP

## Acknowledgements

The authors would like to thank Sashen Moodley, Ashentha Govender and Colin Chetty in the Microbiology Core, Africa Health Research Institute.

