## Supplementary Material for "Whole genome sequencing *Mycobacterium tuberculosis* directly from sputum identifies more genetic diversity than sequencing from culture"


| | |
|---|---|
| Supplementary Methods | Page 2 |
| Supplementary Figures | Page 4 |
| Supplementary Table | Page 7 |

### Supplementary Methods

**Microbiology**
The solid agar proportion method was used to perform phenotypic DST for Durban samples. DST was done for isoniazid (minimum inhibitory concentrations of 0.2µg/ml [low-level resistance] and 1.0µg/ml [high-level resistance]), rifampicin (1.0µg/ml), ethambutol (7.5µg/ml), streptomycin (2.0µg/ml), ofloxacin (2.0µg/ml) and kanamycin (6.0µg/ml).

**Bioinformatic analysis**

Command line parameters used were as follows:

```
trim_galore = Trim Galore v0.4.4
bbmap = BBMap v38.32
picard = Picard Tools v1.13
qualimap = Qualimap v2.21
freebayes = FreeBayes v1.2
vcffilter and vcfintersect from vcflib v1.0
samtools = Samtools v1.9
varscan = VarScan v2.4.0

####Trimming
echo "Trimming"
$trim_galore --length 50 --no_report_file --paired $file1 $file2

####Build BBMap index
$bbmap ref=$reference

###########BBMap align reads
echo "Mapping reads"
$bbmap -Xmx20g in=$file1trim in2=$file2trim out=$sample.aln.sam ref=$reference t=16
minid=0.98 pairedonly=t pairlen=500 slow=t statsfile=$sample'.stats'

########### Sort sam file
echo 'Sorting .sam file to .bam file...'
AlignmentBAM=$sample.sorted.bam
java -Xmx10g -jar $picard SortSam I=$sample.aln.sam O=$AlignmentBAM
SORT_ORDER=coordinate MAX_RECORDS_IN_RAM=1000000

########### Remove duplicates
echo 'Removing duplicates...'
DeduplicatedBAM=$sample.dedup.bam
Metrics=$sample.metrics.txt
java -Xmx10g -jar $picard MarkDuplicates REMOVE_DUPLICATES=true I=$AlignmentBAM
O=$DeduplicatedBAM METRICS_FILE=$Metrics
```

```
############ Add name to read group
mkdir BBMap098_new
DeduplicatedBAM1='BBMap098_new/'$sample'.dedup.1.bam'
java -Xmx10g -jar $picard AddOrReplaceReadGroups LB=BRC PL=illumina PU=Mtb
SM=$sample I=$DeduplicatedBAM O=$DeduplicatedBAM1

############ Sort bam file
java -Xmx10g -jar $picard BuildBamIndex INPUT=$DeduplicatedBAM1

############ Quality checking
mkdir qualimap_098new
$qualimap bamqc -bam $DeduplicatedBAM1 -outdir qualimap_098new/$sample

############# FreeBayes SNP calling, filter mapping quality at 30 and base quality at 30,
filter PPE genes with vcfintersect
echo 'FreeBayes SNP calling...'
mkdir Variants_DOWN_BBMAP098
FreeBayes_permissive_VCF='Variants_DOWN_BBMAP098/'"$sample"'.vcf'
$freebayes -f $reference -p 1 -m 30 -q 30 -C 10 -b
'BAMs_098_DOWN/'$sample'.down.bam'| $vcfintersect -b $PPEgenes -v | $vcffilter -f "SAF
> 0 & SAR > 0 & AO > 3 & RPL > 0 & RPR > 0"> $FreeBayes_permissive_VCF;
$vcfintersect -b $PPEandRNA -v $FreeBayes_permissive_VCF >
$FreeBayes_permissive_VCF'.noRNA'

############# Filter heterozygous alleles
FreeBayes_het_VCF='Variants_DOWN_BBMAP098/'"$sample"'.FreeBayes.het.vcf'
$vcffilter -f "SRF > 0 & SRR > 0 & RO > 3" $FreeBayes_permissive_VCF >
$FreeBayes_het_VCF
$vcfintersect -b $PPEandRNA -v $FreeBayes_het_VCF > $FreeBayes_het_VCF'.noRNA'

############# Calling variants with VarScan for consensus sequence
mkdir Consensus
echo "Making varscan file..."
$samtools mpileup -B -f NC_000962.3.fasta --positions
PE_PPE_RNA_INVERSE_Comas_190212.bed -q 30 -Q 30
'BAMs_098_DOWN/'$sample'.down.bam' \
| java -Xmx8g -jar $varscan mpileup2cns --min-freq-for-hom 0.95 --min-var-freq 0.95 --min-
coverage 20 --p-value 99e-02 --min-avg-qual 30 > 'Consensus /'$sample'.varscanout'

############# Generate consensus of equal length to reference
echo "Making fasta"
perl varscan_to_pseudoseq.pl 4411532 'Consensus/'$sample'.varscanout' >
'Consensus/'$sample'.fasta'
```

### Supplementary Figures

Supplementary Figure 1. Midpoint rooted maximum likelihood phylogenetic tree of all samples. Nodes annotated with bootstrap values. WGS obtained from MGIT are appended _M and those directly from sputum _S.

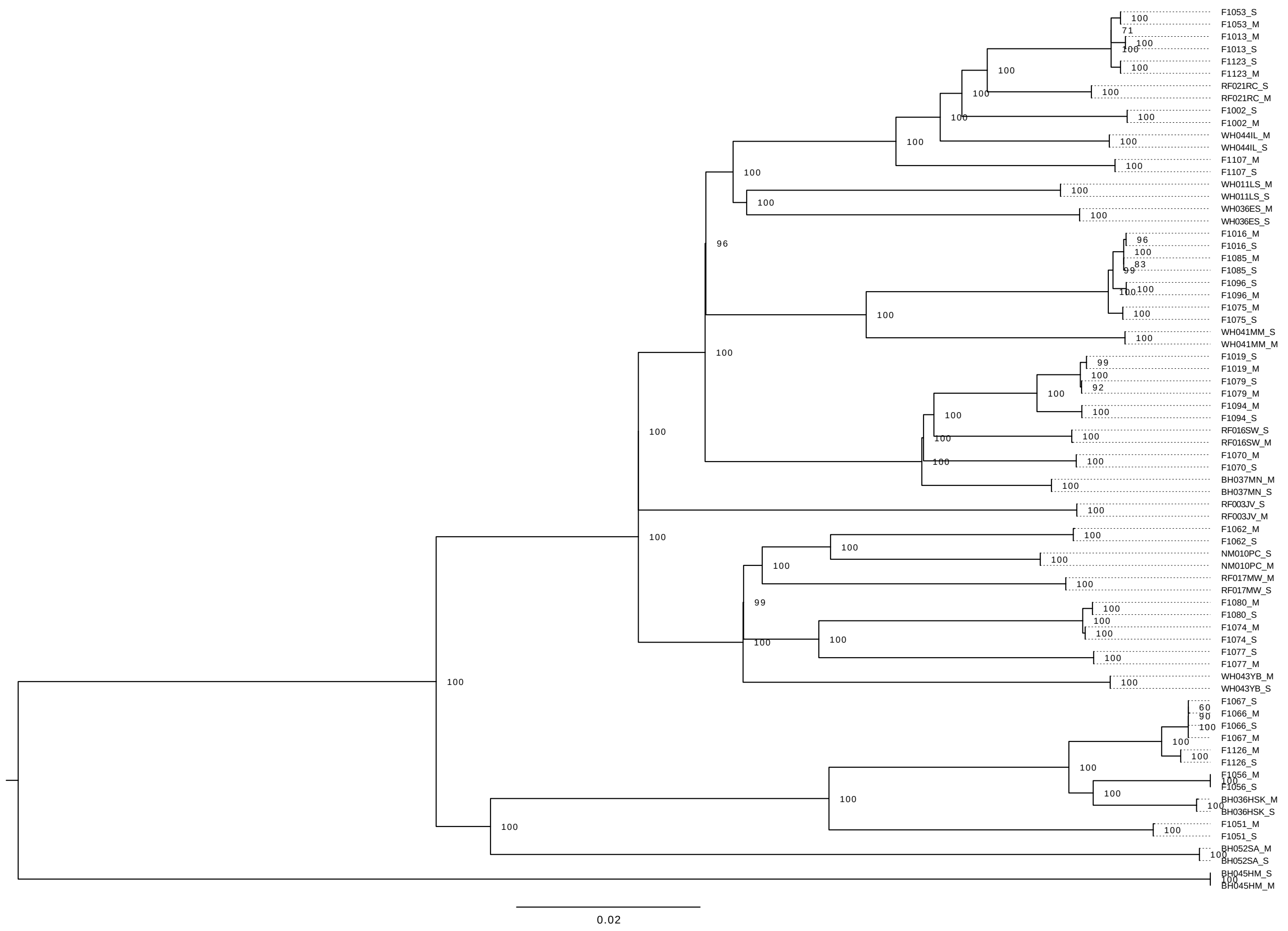

Supplementary Figure 2. Percentage of sampled reads not assigned to *M. tuberculosis* by Kraken (see Methods) that had a blast hit of ≥30 bases to *M. tuberculosis* ribosomal RNA genes (16S or 23S rRNA from H37Rv; see Methods). Bars show median and interquartile range.

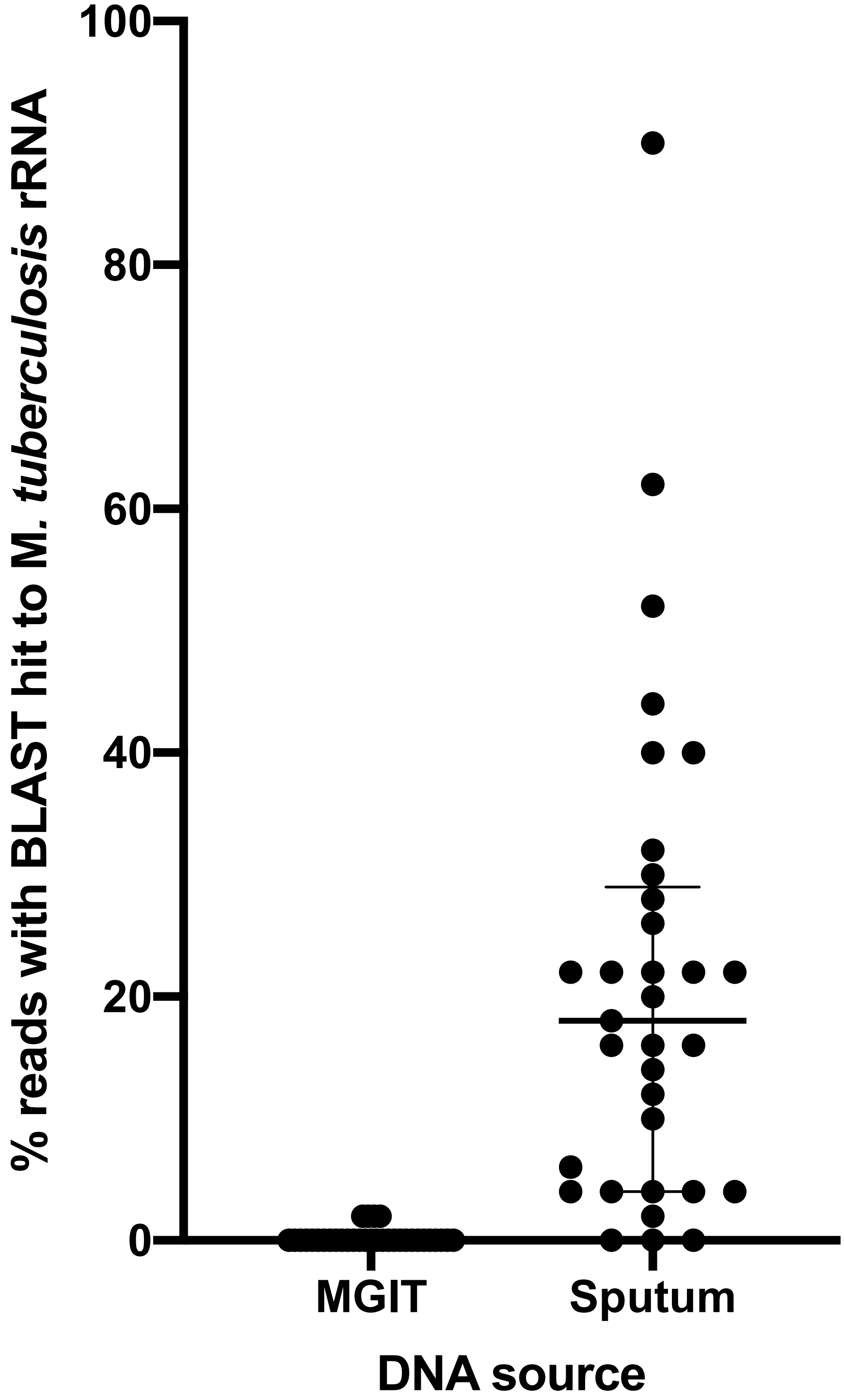

Supplementary Figure 3. Total subsampled reads with a BLAST hit to *M. tuberculosis* ribosomal RNA genes, with colours indicating taxonomic assignment with Kraken (see Methods).

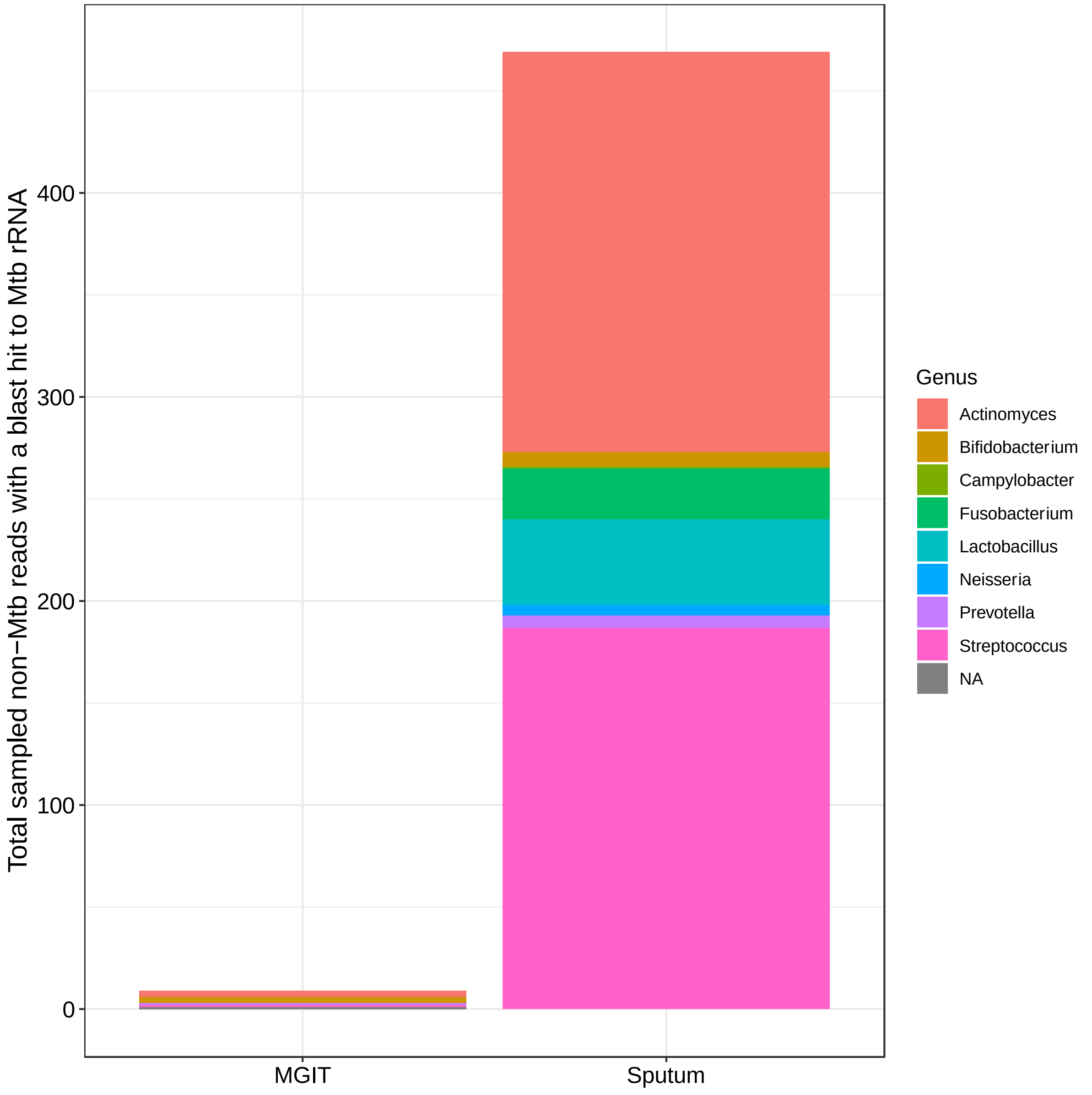

### Supplementary Tables

Supplementary Table 1. Original mean coverage depth and number of heterozygous alleles (HAs) after exclusion of hypervariable (e.g. PE/PPE) and ribosomal RNA genes. Patient F1096 had evidence of mixed infection in the sputum sequence and so was excluded from analysis.

| Patient ID | Mean coverage depth | | % covered at 20x | | HA count | |
|---|---|---|---|---|---|---|
| | MGIT | Sputum | MGIT | Sputum | MGIT | Sputum |
| BH037 | 71.9 | 97.3 | 95.9% | 90.3% | 1 | 2 |
| BH052 | 113.3 | 244.2 | 96.6% | 90.0% | 21 | 11 |
| F1002 | 73.9 | 98.7 | 96.8% | 92.1% | 5 | 6 |
| F1013 | 96.1 | 135.0 | 97.1% | 91.6% | 2 | 4 |
| F1016 | 58.8 | 255.4 | 95.6% | 96.1% | 3 | 4 |
| F1019 | 161.5 | 76.7 | 97.1% | 88.6% | 4 | 6 |
| F1051 | 241.5 | 293.4 | 97.3% | 95.6% | 7 | 5 |
| F1053 | 199.5 | 216.5 | 97.8% | 95.2% | 2 | 4 |
| F1056 | 256.1 | 92.9 | 96.4% | 89.5% | 7 | 12 |
| F1062 | 123.6 | 264.2 | 96.9% | 95.9% | 8 | 5 |
| F1066 | 77.8 | 563.9 | 95.8% | 96.8% | 4 | 5 |
| F1067 | 232.4 | 364.9 | 95.5% | 96.5% | 11 | 7 |
| F1074 | 184.5 | 173.7 | 97.7% | 95.1% | 3 | 4 |
| F1075 | 161.4 | 101.3 | 97.2% | 90.9% | 3 | 4 |
| F1077 | 85.4 | 156.1 | 95.9% | 93.8% | 2 | 4 |
| F1079 | 118.8 | 245.2 | 92.2% | 95.7% | 4 | 5 |
| F1080 | 97.7 | 94.1 | 93.5% | 90.2% | 6 | 9 |
| F1085 | 245.6 | 293.8 | 97.9% | 97.0% | 12 | 10 |
| F1094 | 87.0 | 368.5 | 94.0% | 96.7% | 5 | 5 |
| F1096 | 147.5 | 60.9 | 96.7% | 85.8% | 3 | 329 |
| F1107 | 138.0 | 417.5 | 96.6% | 97.4% | 0 | 1 |
| F1123 | 124.5 | 111.1 | 96.8% | 93.1% | 2 | 5 |
| F1126 | 101.1 | 314.8 | 95.8% | 96.2% | 6 | 5 |
| NM010 | 164.9 | 167.4 | 98.0% | 90.3% | 7 | 7 |
| RF003 | 139.4 | 80.4 | 98.4% | 86.7% | 5 | 5 |
| RF016 | 142.4 | 314.2 | 96.7% | 97.2% | 2 | 1 |
| RF017 | 230.1 | 101.8 | 97.5% | 92.9% | 13 | 13 |
| RF021 | 147.4 | 214.1 | 97.3% | 95.8% | 11 | 23 |
| WH011 | 132.5 | 172.0 | 98.4% | 92.5% | 4 | 4 |
| WH036 | 179.3 | 127.6 | 97.1% | 87.4% | 3 | 20 |
| WH041 | 188.5 | 102.2 | 97.5% | 89.2% | 0 | 8 |
| WH043 | 211.5 | 286.8 | 98.4% | 97.0% | 15 | 16 |
| WH044 | 169.0 | 217.0 | 98.4% | 95.6% | 22 | 45 |

Supplementary Table 2. Intergenic regions with ≥2 heterozygous alleles (HAs) across all sputum samples, ordered by greatest number of HAs per base.

| **Intergenic region** | **HAs per base** | | **Total number of HAs** | |
|---|---|---|---|---|
| | **Sputum** | **MGIT** | **Sputum** | **MGIT** |
| *pe_pgrs18-mprA* | 0.043 | 0.022 | 16 | 8 |
| *pe_pgrs45-rv2616* | 0.041 | 0.029 | 14 | 10 |
| *nrdH-rv3054c* | 0.040 | 0.015 | 19 | 7 |
| *proT-vapC12* | 0.017 | 0.003 | 6 | 1 |
| *alr-rv3424c* | 0.017 | 0.003 | 5 | 1 |
| *pe_pgrs17-rv0979c* | 0.013 | 0.000 | 4 | 0 |
| *rv2355-ppe40* | 0.009 | 0.000 | 7 | 0 |
| *lgt-rv1615* | 0.007 | 0.000 | 5 | 0 |
| *rv0794c-rv0795* | 0.007 | 0.005 | 3 | 2 |
| *rv3428c-ppe59* | 0.006 | 0.000 | 7 | 0 |
| *pe8-rv1041c* | 0.002 | 0.001 | 2 | 1 |

Supplementary Table 3. National Center for Biotechnology Information Sequence Read Archive (NCBI SRA) accession numbers for each sample.

| Patient ID | NCBI SRA Accession Number | |
|---|---|---|
| | MGIT | Sputum |
| BH037 | SRR7725437 | SRR7725438 |
| BH052 | SRR7725441 | SRR7725442 |
| F1002 | SRR7725443 | SRR7725444 |
| F1013 | SRR7725410 | SRR7725411 |
| F1016 | SRR7725404 | SRR7725405 |
| F1019 | SRR7725406 | SRR7725407 |
| F1051 | SRR7725416 | SRR7725417 |
| F1053 | SRR7725368 | SRR7725367 |
| F1056 | SRR7725370 | SRR7725369 |
| F1062 | SRR7725374 | SRR7725373 |
| F1066 | SRR7725376 | SRR7725375 |
| F1067 | SRR7725383 | SRR7725389 |
| F1074 | SRR7725397 | SRR7725398 |
| F1075 | SRR7725391 | SRR7725395 |
| F1077 | SRR7725414 | SRR7725415 |
| F1079 | SRR7725390 | SRR7725412 |
| F1080 | SRR7725388 | SRR7725387 |
| F1085 | SRR7725386 | SRR7725385 |
| F1094 | SRR7725384 | SRR7725432 |
| F1096 | SRR7725393 | SRR7725392 |
| F1107 | SRR7725420 | SRR7725421 |
| F1123 | SRR7725422 | SRR7725423 |
| F1126 | SRR7725424 | SRR7725425 |
| NM010 | SRR7725426 | SRR7725427 |
| RF003 | SRR7725428 | SRR7725429 |
| RF016 | SRR7725380 | SRR7725379 |
| RF017 | SRR7725382 | SRR7725381 |
| RF021 | SRR7725402 | SRR7725400 |
| WH011 | SRR7725378 | SRR7725433 |
| WH036 | SRR7725418 | SRR7725419 |
| WH041 | SRR7725403 | SRR7725394 |
| WH043 | SRR7725401 | SRR7725396 |
| WH044 | SRR7725430 | SRR7725413 |
